# A discrete character evolution model for phylogenetic comparative biology with Γ-distributed rate heterogeneity among branches of the tree

**DOI:** 10.1101/2024.05.25.595896

**Authors:** Liam J. Revell, Luke J. Harmon

**Affiliations:** Department of Biology, University of Massachusetts Boston, Boston, MA, USA; Facultad de Ciencias, Universidad Católica de la Santísima Concepción, Concepción, Chile; Department of Biological Sciences, Institute for Bioinformatics and Evolutionary Studies (IBEST), University of Idaho, Moscow, ID, USA

**Author notes:** Corresponding author: Liam J. Revell^1,2^.

## Abstract

Phylogenetic comparative methods are now widely used to measure trait evolution on the tree of life. Often these methods involve fitting an explicit model of character evolution to trait data and then comparing the explanatory power of this model to alternative scenarios. In this article, we present a new model for discrete trait evolution in which the rate of character change in the tree varies from edge (i.e., “branch”) to edge of the phylogeny according to a discretized Γ distribution. When the edge-wise rates of evolution are, in fact, Γ-distributed, we show via simulation that this model can be used to reliably estimate the shape parameter (*α*) of the distribution of rate variation among edges. We also describe how our model can be employed in ancestral state reconstruction, and demonstrate via simulation how doing so will tend to increase the accuracy of our estimated states when the generating edge rates are Γ-distributed. We discuss how marginal edge rates are estimated under the model, and apply our method to a real dataset of digit number in squamate reptiles, modified from Brandley et al. (2008).

## 1 INTRODUCTION

A common endeavor in phylogenetic comparative analysis combines an estimated phylogeny with species trait data in an effort to better understand the evolution and history of our study clade (Harmon 2019; Revell and Harmon 2022). So doing frequently requires that we specify an explicit mathematical model that postulates how our traits, such as body size, limb lengths, the presence or absence of some feature, or a category of habitat specialization, may have evolved on the tree over macroevolutionary time (Pagel 1994; Butler and King 2004; O’Meara et al. 2006; O’Meara 2012; Beaulieu et al. 2013; Harmon 2019). This model might take a variety of flavors, all with its own parameters and attributes. In this case, our analysis may simply involve fitting a set of such models, each capturing a slightly different scenario of trait evolution, to learn which of them best explains the data that we have in our possession (e.g., Revell et al. 2022). Once we’ve identified a suitable trait evolution model, we might use it to estimate model parameters or reconstruct ancestral states (Harmon 2019; Revell et al. 2022; Revell 2024).

When our trait data are discretely-valued (that is, the character only manifests in one of a limited set of conditions), the predominant model employed by comparative biologists to approximate its evolution on the tree is one that’s popularly referred to as the M*k* model (Pagel 1999; Lewis 2001; Harmon 2019; Revell and Harmon 2022). This model is so named because the underlying stochastic process it postulates is a memoryless Markov chain (the ‘M’), with a *k*-dimensional state space (Lewis 2001). What we refer to here as the ‘state space’ is just the number of unique conditions that the Markov chain can assume: two, for example, if our character can take on two different levels; three, if it might exist in three values; etc. Though originally conceived as a model with equal back-and-forth rates of transition between the two, three, or more conditions in the space (Lewis 2001), the M*k* model has now been elaborated to permit an arbitrary number of different rates of character transition between levels (described in Harmon 2019; Revell and Harmon 2022). Fitting such a model could involve estimating up to *k ×* (*k* − 1) parameters for *k* states (all non-diagonal elements of a *k ×k* state transition matrix, typically called the **Q** matrix).

Many of our comparative methods for analyzing trait change assume a uniform tempo of evolutionary change across all the branches and nodes of the tree (Harmon 2019). This is despite the fact that empirical results strongly suggest that the rate of change in phenotypic traits can vary widely across lineages and through time (e.g., Mahler et al. 2010; Beaulieu et al. 2013; Sander et al. 2021). Even if we are not interested in rate heterogeneity as such, to the extent that we are using a fitted trait model for other inferences, such as ancestral state reconstruction, we should worry that rate heterogeneity can lead us astray. For example, consider the situation where a trait changes only rarely, even over deep time – but also exhibits rapid change within certain young clades, perhaps for some particular ecological or developmental reason. Imagine that a scientist studying this trait is interested in ancestral character conditions in the clade and fits a homogeneous rate model: since this is (by far) the most common assumption of ancestral character evolution. A homogeneous model with a low rate of change between states will do a poor job explaining rapid evolution of the trait where it occurred, even if it could do a good job explaining our data across the rest of our phylogenetic tree. As a consequence, our fitted model will have a high overall evolutionary rate which, when applied across the tree, greatly increases uncertainty of all ancestral states throughout the phylogenetic history of our group.

The M*k* model has already been extended in a number of interesting ways to permit rate heterogeneity over time, among clades, or as a function of another trait that has been mapped along the nodes and edges of the phylogeny. For instance, Pagel (1994) described a clever manner in which an M*k*-type model can be used to approximate the interdependent evolution of two different binary traits, in which the condition of character one influence the transition rates of character two and/or vice versa. Later, Marazzi et al. (2012) and Beaulieu et al. (2013) proposed a different model (called the precursor or hidden-rates model, depending on the way in which it’s parameterized) whereby one or multiple unobserved conditions can influence our trait’s transition rate to other observed values (also see Revell and Harmon 2022). Revell and Harmon (2022) described a polymorphic trait evolution model in which the polymorphic condition (e.g., *a* + *b*) is postulated to be evolutionarily intermediate between corresponding monomorphic states (*a* and *b*, in this case). Revell et al. (2021) defined a model in which the rate of evolution of a discrete character is influenced by an explicit, *a priori* hypothesis of rate heterogeneity that’s been painted onto the edges and nodes of the phylogeny by the investigator. Lastly, various relaxed molecular clock models have been described in the literature (e.g., Thorne et al. 1998; Huelsenbeck et al. 2000; Yoder and Yang 2000; Drummond and Suchard 2010), and these models are sometimes used to study variable-rate phenotypic trait evolution on trees (e.g., King and Lee 2015).

In the present article, we’ll describe a new extension of the M*k* model in which the edge-wise rates of evolution of our trait are sampled randomly from a (normalized) Γ distribution with shape parameter *α*. This model is in many ways analogous to the popular Γ rate heterogeneity model for molecular sequence evolution that’s commonly used for phylogeny estimation (e.g., Yang 1994; Felsenstein 2004). In our case, however, we’ll be assuming that that rate of evolution for a single discrete trait changes from edge to edge according to an uncorrelated Γ (rather than varying by nucleotide site). This article is intended to provide a description of the method, which has already been implemented in the popular R package *phytools* (Revell 2012; Revell 2024), a preliminary exploration of its statistical properties, and an empirical example based on the evolution of digit number in squamate reptiles using data adapted from Brandley et al. (2008).

## 2 MODEL, METHODS, AND RESULTS

### 2.1 The model

This article describes a new discrete phenotypic trait evolution model in which the relative rates of character evolution varies from edge to edge according to an uncorrelated Γ distribution with shape parameter *α* and mean of 1.0. (A mean of 1.0 is obtained by setting the second parameter of the Γ distribution, *β*, to *β* = *α*.) Note that, much as it is for models of molecular sequence evolution, this assumption does not constrain the average rate of character change, nor the relative rates of state changes of different types – both of which are given by the transition matrix of the process, **Q**.

The Γ distribution is a flexible probability distribution that takes a shape resembling that of an exponential function for low values of *α* (e.g., *α* ≤ 1.0), and an increasingly bell-like conformation for larger and larger values of *α* (e.g., *α* ≥ 4). Figure 1A shows a set of Γ distributions for various values of *α*.

**Figure 1.**
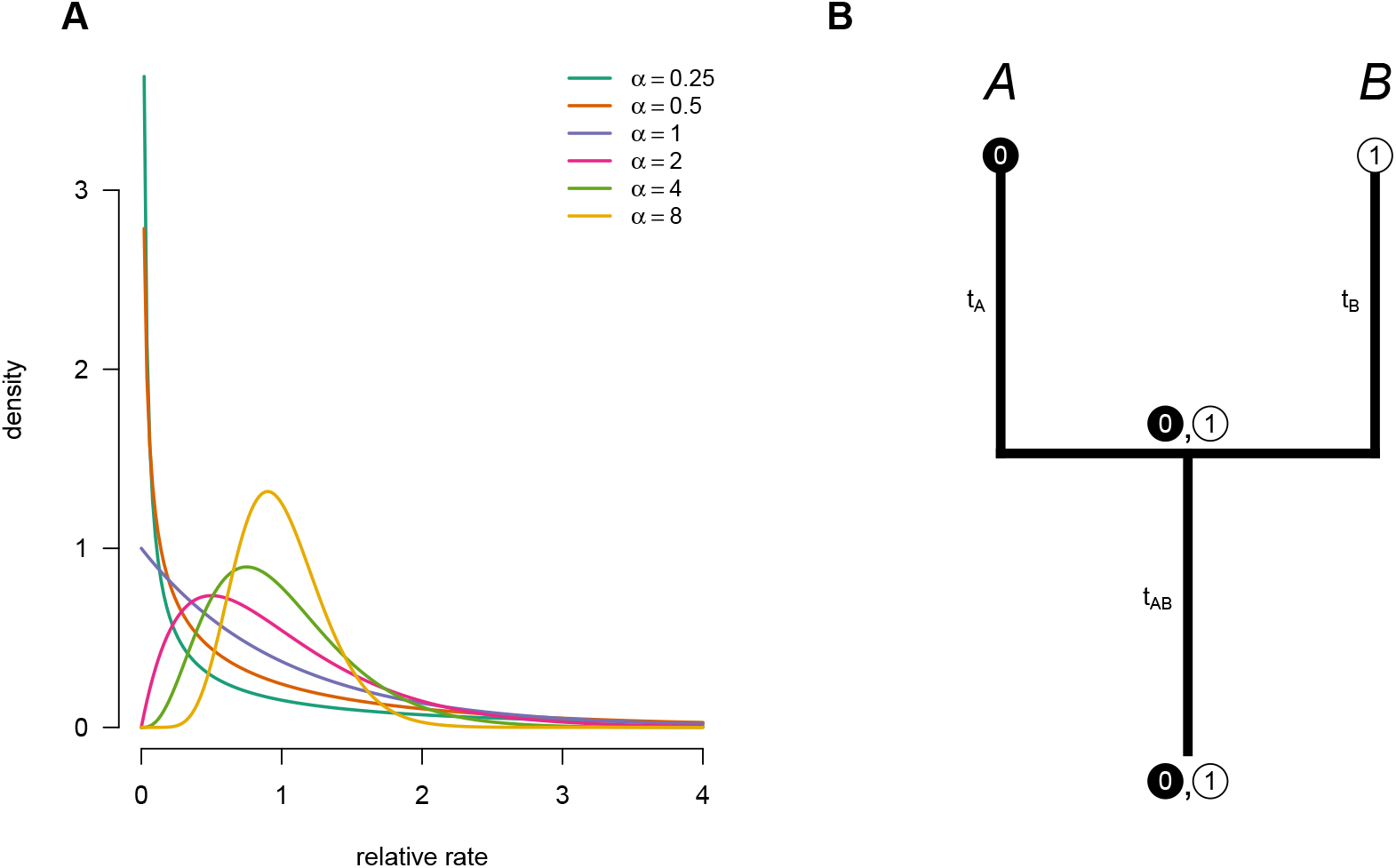
(**A**) A set of Γ probability distribution with different values of the shape parameter, *α*. By setting *β* = *α* the mean of each distribution is equal to 1.0, while *α* determines the shape of the distribution in which *α <* 1 correspond to distributions with high among edge rate heterogeneity. (**B**) An example phylogeny with two taxa (*A* and *B*) and two trait values of a discrete character (0 and 1). See main text for more details.

Under a standard M*k* model, the rates of transition between character levels are dictated entirely by a *k ×k* matrix **Q** in which each element, *q*_*i, j*_ (for all *i≠ j*), gives the instantaneous transition rate from condition *i* to condition *j*, and the diagonal is equal to the negative row sum (Revell and Harmon 2022). For a binary trait (i.e., *k* = 2), this transition matrix (**Q**) would appear as follows.

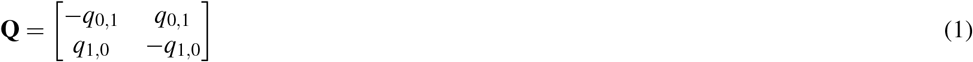

To compute the probability of character change under a standard M*k* model, given **Q** and any arbitrary elapsed time, *t*, we simply calculate **P** = exp(**Q** *×t*), in which exp(**X**) indicates the matrix exponential of **X**. This operation gives us a new *k ×k* matrix, **P**, in which *p*_*i, j*_ is equal to the probability of beginning the time interval, *t*, in condition *i* and ending in condition *j* under our modeled process.

To see how we then go about calculating the probability of a set of data at the tips of the tree for any particular value of **Q**, we can begin by considering the simplified tree and data pattern of Figure 1B. In this tree, the two nominal taxa of our phylogeny, *A* and *B*, each exhibit different conditions for the character: taxon *A* has the state 0 (shown in black); while taxon *B* is found in condition 1 (shown in white). Logically, the probability of observing this set of states must be equal to the probability of starting (at the root) in state 0, here written as *π*_0_, multiplied by the probability of finding the single internal node of our tree in state 0 after elapsed time *t*_*AB*_, *P*(0|0, *t*_*AB*_), multiplied by the probability of the tip *A* being observed in state 0 after elapsed time *t*_*A*_, *P*(0|0, *t*_*A*_), multiplied in turn by the probability of tip *B* being observed in state 1 after elapsed time *t*_*B*_, *P*(1|0, *t*_*B*_) – and then summed across both possible root state and all four possible internal node state combinations. When written down, this probability is as follows.

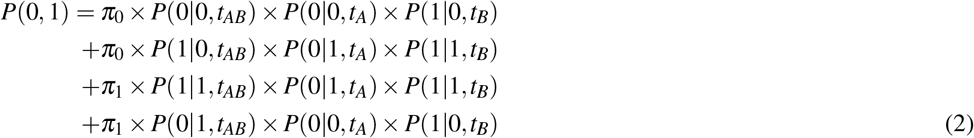

Though for a two-taxon tree, such as the one shown in Figure 1B, the value of equation (2) is straightforward to compute, doing so in the same way for a tree with three, four, or more species would be increasingly onerous if it involved the explicit enumeration of all possible states at all internal nodes of the phylogeny, as we’ve done here. Fortunately, Felsenstein (1981) described an efficient “pruning” algorithm to compute this probability via a single post-order (tip to root) traversal of the tree.

Now to account for variation in the rate of evolution from edge to edge of our phylogeny, we’ll use a discretization of the Γ distribution following Yang (1994). Directly following Yang (1994), we first set *m* rate categories (*k* in Yang 1994, but *m* here to avoid confusion with our *k* different character states). We then subdivide the range of our relative rates, *r* (which could theoretically vary from 0 to ∞, the range of a Γ-distributed random variable), into *m* evenly spaced bins. To compute the median of each bin, we need to find the values of each *r*_*i*_ (that is, *r*_1_, *r*_2_, …, *r*_*m*_) at the percentiles of our Γ distribution that correspond to the midpoints of each of *m* bins: 1*/*(2*m*), 3*/*(2*m*), …, (2*m*− 1)*/*(2*m*) (Yang 1994): quantities that are straightforward to calculate using a computer. Referring back, once again, to our two-taxon case of Figure 1B, the probability of our data in which the relative edge rates vary as a random variable drawn from a discretized, uncorrelated Γ distribution can be approximated as follows, in which *r*_*A*_ = *r*_*i*_ indicates that the edge leading to taxon *A* has relative rate *r*_*i*_, and so on.

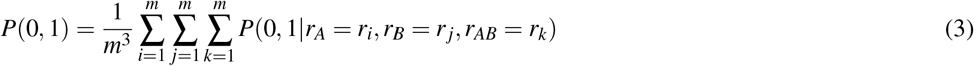

Figure 2 illustrates a complete enumeration of the 3^3^ = 27 possible ways that *m* = 3 rates could be distributed across the three edges of our tree of Figure 1B, along with the probability of our observed data at the tips of the tree for *q*_0,1_ = *q*_1,0_ = 0.1 and *α* = 0.7. To calculate the total probability under this model, we would just sum all of these quantities and divide by 27, as given in equation (3), above.

**Figure 2.**
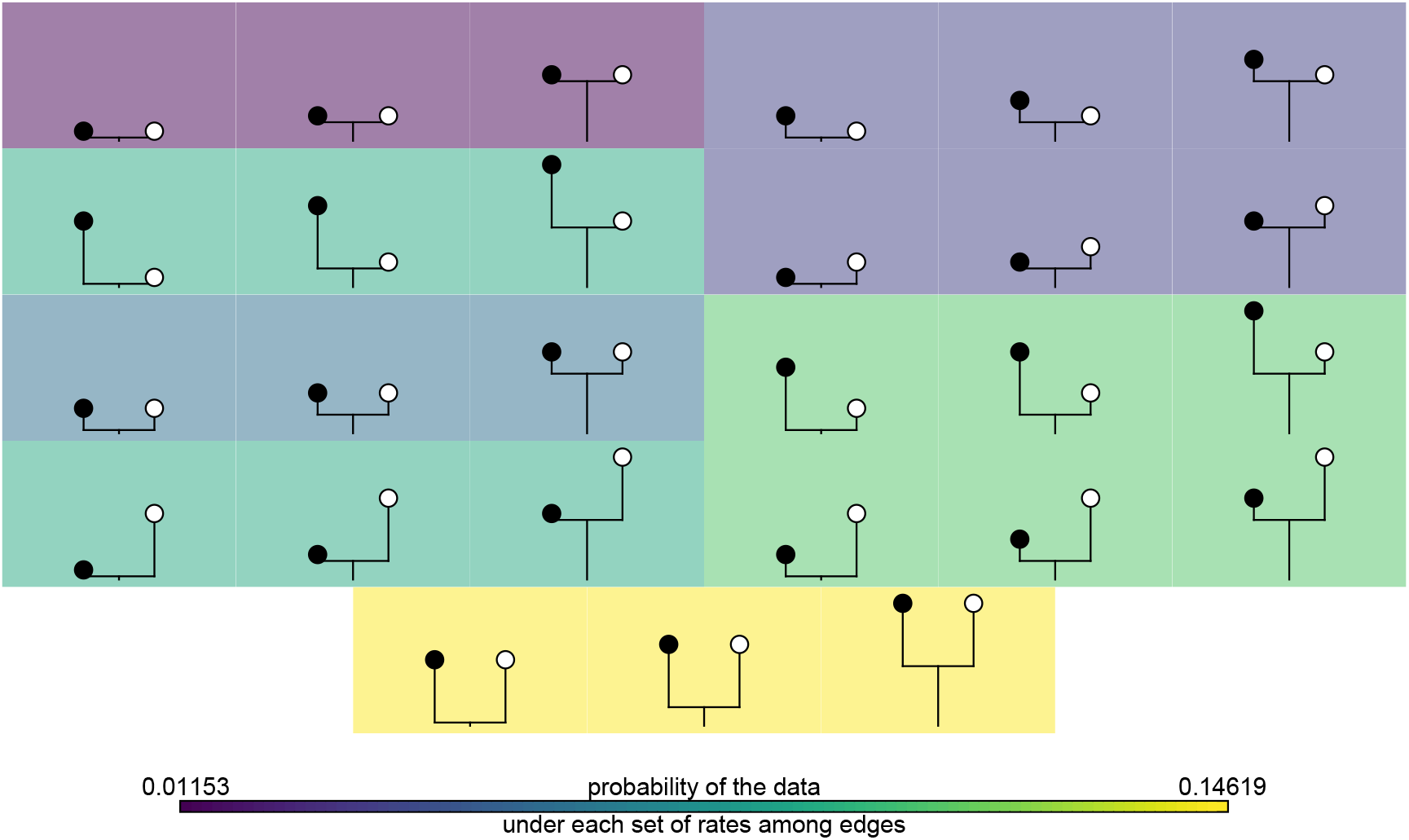
The probability of the tip data from Figure 1B given all the 27 possible ways in which three rate categories (i.e., *m* = 3) could be distributed across the three edges of our two taxon tree, in which *q*_0,1_ = *q*_1,0_ = 0.1 and *α* = 0.7. To compute the total probability of the data at the tips of the phylogeny in Figure 1B whilst integrating over discretized Γ-distributed rates across edges, we can simply sum these values and divide by 27.

As before, this sum would be very onerous to compute (and, indeed, impossible for even relatively modest-sized trees) were it not for the pruning algorithm of Felsenstein (1981). To implement pruning with (discrete) Γ-distributed rate heterogeneity, we also depend on the further equality that for evolution along a single edge of the tree of length *t*, the probability of ending in state 1 given that the edge began in state 0 can be written as follows.

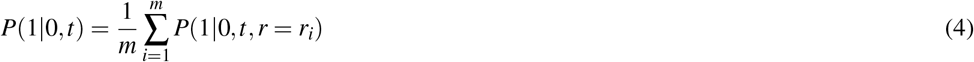

That is to say, the probability of starting in condition 0 and then after some time *t* being in state 1, given that the rate of evolution over time *t* is a discretized Γ-distributed random variable with conditions *r* = *r*_1_, *r*_2_, …, *r*_*m*_ each with probability 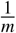, is just the probability of each rate category (here they are even, hence 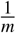) times the probability of the state given that rate. More generally, the *matrix* of transition probabilities between states after time *t* can be expressed as a function of probability of each rate category times the exponentiated product **Q***r*_*i*_*t*, in which exp(**X**) is the matrix exponential of **X**.

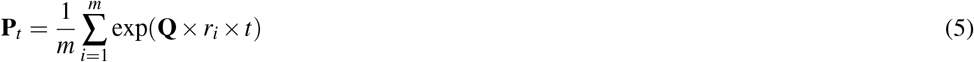

As we can compute the probability of any data pattern on the tree for a given shape parameter (*α*), number of rate categories (*m*), and transition matrix **Q**, we can likewise find the set of such values that maximize this probability – in other words, the Maximum Likelihood (ML) estimate. That’s what we will do here. As a side note to this observation, we’ll also take this opportunity to observe that it would be relatively straightforward to fit the model of this article using Bayesian MCMC instead of ML. Indeed, the *phytools* function in which this model has been implemented (`fitgammaMk`), also exports the likelihood expression of the model that *phytools* users could easily pass along their favorite MCMC algorithm. (This would, of course, require the investigator to supply appropriate prior probability distributions for *α* and **Q**, a task that has been contemplated in other literature.)

In this article, we focus on three ways that readers might consider making use of this model, although others are also certainly possible. First, researchers can estimate *α*, which gives insight into how variable the tempo of evolution has been across the branches or clades of our phylogeny. Low values of *α* correspond to high edge-wise rate heterogeneity, whereas high values of *α*, relatively little rate variation from branch to branch of the tree. Second, investigators can calculate (conditioning on a transition model and ML value for *α*) the marginal likelihoods that each edge of the tree is in each of the *m* rate categories of the fitted model. This will be done simply by traversing the tree, edge by edge, holding the rate on a given edge constant to each of the *m* rate categories, and then computing the marginal probability of the data (integrating over all *m* rates across all of the *other* edges of the tree) for each rate. Since these marginal likelihoods can also be interpreted as the posterior probabilities that each edge is in each of our *m* rate conditions, a researcher can in turn proceed to use them as weights for the computation of marginal edge rates (as the simple weighted mean of the median rate of each category) for each branch of the phylogeny. This will help researchers to measure precisely how variation in the tempo of evolution is distributed across their phylogenetic tree. Finally, third, investigators might use the model to perform marginal or joint ancestral state reconstruction at the nodes of their tree. This is undertaken in exactly the usual way (see Yang 2014; Revell and Harmon 2022), but in which we integrate over each of the *m* rate categories as we traverse the tree. We’ll illustrate all three of these applications below.

### 2.2 Estimation of *α*

Now that we’ve expressed, in principle, how one might go about computing the probability of a set of data for any particular value of the (discretized) Γ distribution shape parameter, *α*, and state transition matrix, **Q**, we used numerical simulations to establish whether or not we can, in fact, estimate *α* in practice. Note that these initial simulations have not been designed to explore the range of conditions (in taxon number, transition models, character state levels, or values of *α*) that might be encountered in an empirical study. To the contrary, we decided to use a small number of ‘maximally-powered’ simulations that we anticipated might give us the *best* chance of reliably measuring rate heterogeneity as a proof-of-concept that Γ-distributed rate heterogeneity of a single discrete phenotypic trait can be detected if it exists. We’ll broaden our simulation conditions in subsequent sections of this article (focused on statistical power and marginal ancestral state estimation), and also hope to see the wider variety of numerical simulations that will be required to understand the study of rate heterogeneity from a single character in future articles by us or others.

Thus, to (as much as reasonably possible) *maximize* our power to estimate *α*, we simulated a total of twenty 2,000 species, pure-birth (Yule) phylogenies, rescaling each tree to have a total depth of 10 time unit. We sampled the relative edge rates of each tree from a Γ distribution with a total of *six* different values of the Γ shape parameter, *α*: 0.25, 0.5, 1.0, 2.0, 4.0, and 8.0. (Note that although our model for estimation is a discretized Γ distribution, we intentionally elected to use a continuous Γ distribution for simulation both here and in subsequent sections of this article. We’re thus also testing how reasonably-well our discretization of Γ approximates underlying continuous rate variation among edges.) These simulated values of *α* varied from very high rate heterogeneity among edges (*α* ≤ 1.0) to relatively low rate heterogeneity (*α* ≥ 4.0; Figure 1A). Having sampled edge rates, we next simulated a single phenotypic character vector on each tree. We used the following generating value of **Q** for an ordered, three-state character with conditions *a, b*, and *c*, and a single transition rate, *q* = 0.5, between adjacent states.

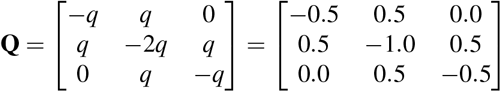

For each tree and data vector, we then proceeded to fit our variable-rate model using ML, specifying the known *structure* of the transition process (that is, the order of character evolution and the number of distinct rate parameters of **Q**), but simultaneously estimating the instantaneous transition rate (*q*) and Γ shape parameter (*α*) – arbitrarily fixing the number of rate categories (*m*) to *m* = 8.

In addition to fitting our Γ rate heterogeneous model, for each tree and dataset we *also* fit a rate homogeneous (i.e., ‘standard’) M*k* model, likewise fixing the model structure to its generating value. We then measured the difference in log-likelihood between the generating rate heterogeneous (Γ) model and our fitted rate homogeneous (M*k*) model. For each phylogeny and set of simulated data, we undertook a total of ten optimization iterations of both the Γ and standard M*k* models, each with random starting values for *α* (if applicable) and the transition rate between states (*q*), to help ensure convergence to the ML solution in each case. We invariably assumed a prior distribution at the root (*π*) following FitzJohn et al. (2009; this has been described as treating the root state as a ‘nuisance parameter’ of the model).

Figure 3A shows the distribution of ML estimated values of *α* for each simulated level of rate heterogeneity. As a general rule, for relatively low values of *α* (that is, high rate heterogeneity among edges: *α* ≤ 1.0), the mean and median estimated values of *α* were quite close to their corresponding generating quantities: highlighted by the horizontal dashed blue lines (Figure 3A). For increasing generating values of *α* (that is, *decreasing* rate heterogeneity among edges), estimated *α* tended to be upwardly biased. This pattern was especially marked for *α* = 8.0, in which many optimization iterations hit the pre-specified upper-bound of optimization (*α* = 1, 000; Figure 3A).

**Figure 3.**
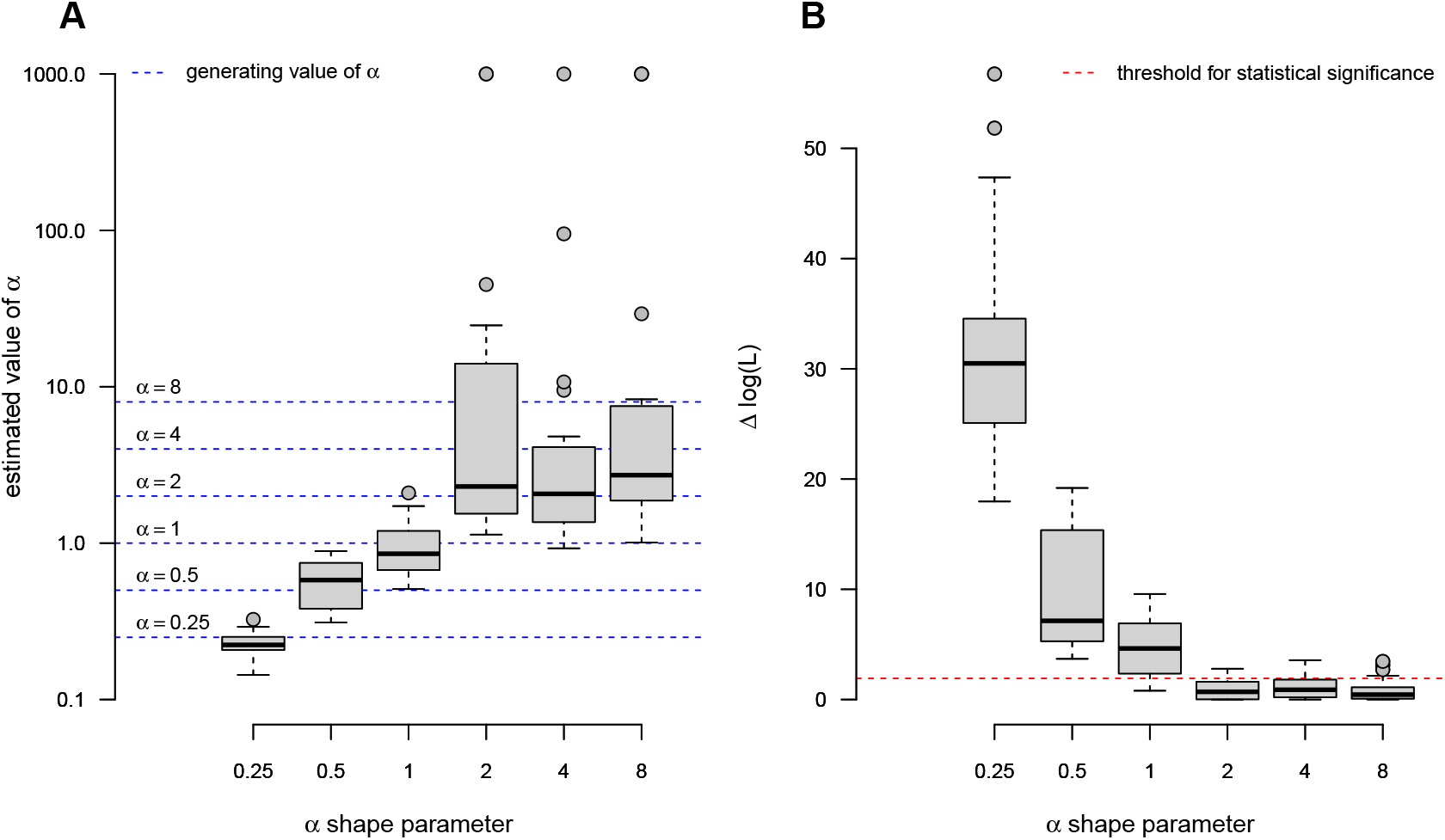
Results from an analysis of simulated datasets generated with Γ-distributed rate heterogeneity under various values of *α*. (**A**) The distribution of estimated values of *α* for each generating value. (**B**) The difference in log-likelihood between the best-fitting Γ model and the standard M*k* rate-homogeneous model. In both panels, the grey box gives the 25% to 75% inter-quartile range, the horizontal black line the median, and the whiskers the range, excluding outliers (shown with grey dots). The maximum upper-bound for *α* during optimization was set to 1,000.

Measurement of the difference in model fit between the Γ rate heterogeneous model and the M*k* model showed a largely consistent trend (Figure 3B). In particular, low *α* (that is, *high* rate heterogeneity among edges) tended to result in much higher model fit of the variable-rate Γ model compared to the standard M*k* model, whereas this advantage declined for increasing *α* (decreasing rate heterogeneity among edges). Notably, even on a tree with 2,000 tips, none of 20 simulation replicates met Δlog-likelihood criterion for statistical significance at the 0.05 level when generating *α* was equal to *α* = 8.0: the lowest level of rate heterogeneity simulated (Figure 3B).

### 2.3 Power analysis

To measure power in a statistical test for significant evidence of rate heterogeneity, we conducted the following simulation. First, we simulated 100 stochastic phylogenies under a Yule (pure-birth) model containing each of 50, 100, 200, 400, 800, and 1,600 species (600 simulations in total), and then rescaled each tree to have a total root to tip depth of 1.0. Next, on each tree we simulated a single discrete character with Γ-distributed rate heterogeneity among edges using an *α* shape parameter of *α* = 0.5, along with the same three-state **Q** matrix as in our prior analysis (see above).

For each simulated tree and data vector, we then proceeded to fit our variable-rate model using ML, specifying the known structure of the model, as before, but simultaneously estimating the instantaneous transition rate (*q*) and shape parameter of Γ, while setting the number of rate categories (*m*) to *m* = 8. In addition to fitting our Γ rate heterogeneous model, for each tree and dataset we also fit a rate homogenous M*k* model, likewise fixing the model structure to its generating value. We then computed a P-value based on a likelihood-ratio test with one degree of freedom: the difference in the number of estimated parameters between our two models. As before, to help ensure convergence to the ML solution we invariably undertook a total of ten optimization iterations of both the Γ and standard M*k* models, each with random starting values for *α* (if applicable) and the transition rate between states (*q*). Finally, we assumed a prior distribution at the root (*π*) following FitzJohn et al. (2009).

The results from our power analysis are shown in Figure 4. We found that the median estimated value of *α* tended to be centered relatively close to the generating value of *α* = 0.5 across all simulation conditions (Figure 4A). The variance among simulations, however, was *very* high for relatively small trees: such as *N* = 50 and *N* = 100 (Figure 4A). Commensurate with this observation, we found that power to reject the null hypothesis of rate homogeneity tended to be barely above the nominal *α* (acceptable type I error) level of 0.05 for small trees, and only increased to around 80% for simulated phylogenies containing as many as 1,600 tip taxa (Figure 4B).

**Figure 4.**
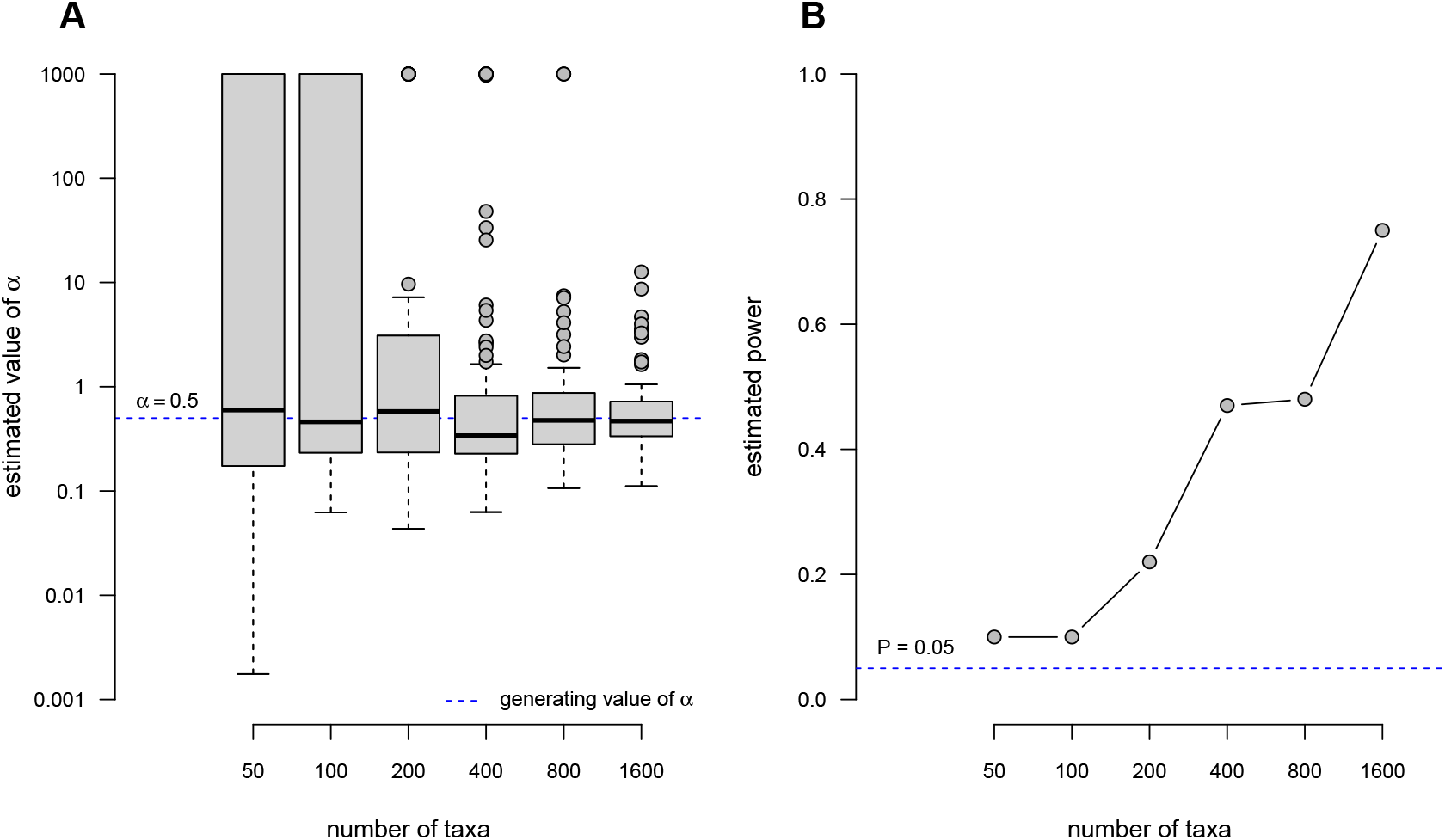
Results from an analysis of simulated datasets generated with Γ-distributed rate heterogeneity with *α* = 0.5 for phylogenies containing various numbers of taxa. (**A**) The distribution of estimated values of *α* for each tree size. As in Figure 3, the grey box gives the 25% to 75% inter-quartile range, the horizontal black line the median, and the whiskers the range, excluding outliers.(**B**) Estimated power: the proportion of statistical test in which Γ rate heterogeneity was significant in a likelihood-ratio test with one degree of freedom, the difference in parameter complexity between the M*k* and Γ models.

### 2.4 Ancestral state reconstruction

Lastly, we measured the accuracy of ancestral state estimation under the Γ model of rate heterogeneity among edges. To do this, we first simulated two hundred 500 taxon pure-birth (Yule) phylogenetic trees, each scaled to a total depth of 1.0. As for prior sections, on each tree we simulated the evolution of a single, three-state discrete character evolving under the same stochastic process (with Γ rate heterogeneity *α* = 0.5) and value of **Q** as in our power analysis of the prior section, but in which we also recorded the simulated ancestral states for all internal nodes of the phylogeny.

Next, for each simulated tree and data vector we proceeded to fit our variable-rate model using ML, specifying the known structure of the model (as in our other simulation analyses), but simultaneously estimating the instantaneous transition rate (*q*) and *α* shape parameter of Γ. As before, we fixed the number of rate categories of our discrete Γ distribution (*m*) to be *m* = 8. In addition to fitting our Γ rate heterogeneous model, for each tree and dataset we also fit a rate homogenous M*k* model, likewise fixing the model structure to its generating value. As before, we undertook a total of ten optimization iterations of both the Γ and standard M*k* models, each with random starting values for *α* (if applicable) and the transition rate between states (*q*), to help ensure convergence. As before, we assumed a FitzJohn et al. (2009) root node prior distribution.

For each fitted M*k* and Γ model of each simulated tree and dataset, we undertook marginal ancestral state estimation following Pagel (1999; Yang 2006). Marginal ancestral state estimation involves traversing the tree by node, and at each node computing the probability of the data given each state of the character, conditioning on the tree and fitted character model, but integrating across all possible states at all other nodes of the tree. These node likelihoods are then normalized (scaled) to sum to 1.0, at which point they can be properly interpreted as the (empirical Bayes) posterior probabilities that each node is in each of the observed conditions of our character (Yang 2006, 2014). To measure the accuracy of our ancestral state reconstructions under both the standard M*k* and Γ models, we simply computed the average, node-wise product of the known, simulated states (represented as vectors of 0s and 1s) and the marginal scaled likelihoods from our procedure of ancestral state estimation. This average product can range from 0 (if all nodes are estimated confidently but completely inaccurately) to 1 (reflecting high confidence in the correct ancestral conditions across all nodes). Intermediate measured accuracy comprises both the scenario of an intermediate portion of nodes inferred correctly with high confidence (and the remainder confidently wrongly), or equally low confidence across all nodes. Since here we’re focused on identifying any difference in accuracy between M*k* and Γ models that might arise for data generated under a Γ model of rate heterogeneity, we’re not too preoccupied in differentiating these latter possibilities. Lastly, we computed the difference in raw accuracy and divided it by the accuracy of the M*k* estimates to measure the proportional improvement in accuracy of the Γ compared to M*k* model when the former was true. (A positive value of 0.05, for example, would indicate that measured accuracy was about 5% better for the Γ model than the M*k* model, whereas a value of −0.05 would suggest the converse.)

Figure 5 shows the distribution of scaled differences in accuracy between the Γ and M*k* models in marginal ancestral state estimation from our 200 simulations. Invariably, estimates obtained using the Γ model, which was the generating evolutionary scenario of the simulations, were more accurate than those obtained from the M*k* model (Figure 5), with a mean improvement in accuracy of about 5.3%.

**Figure 5.**
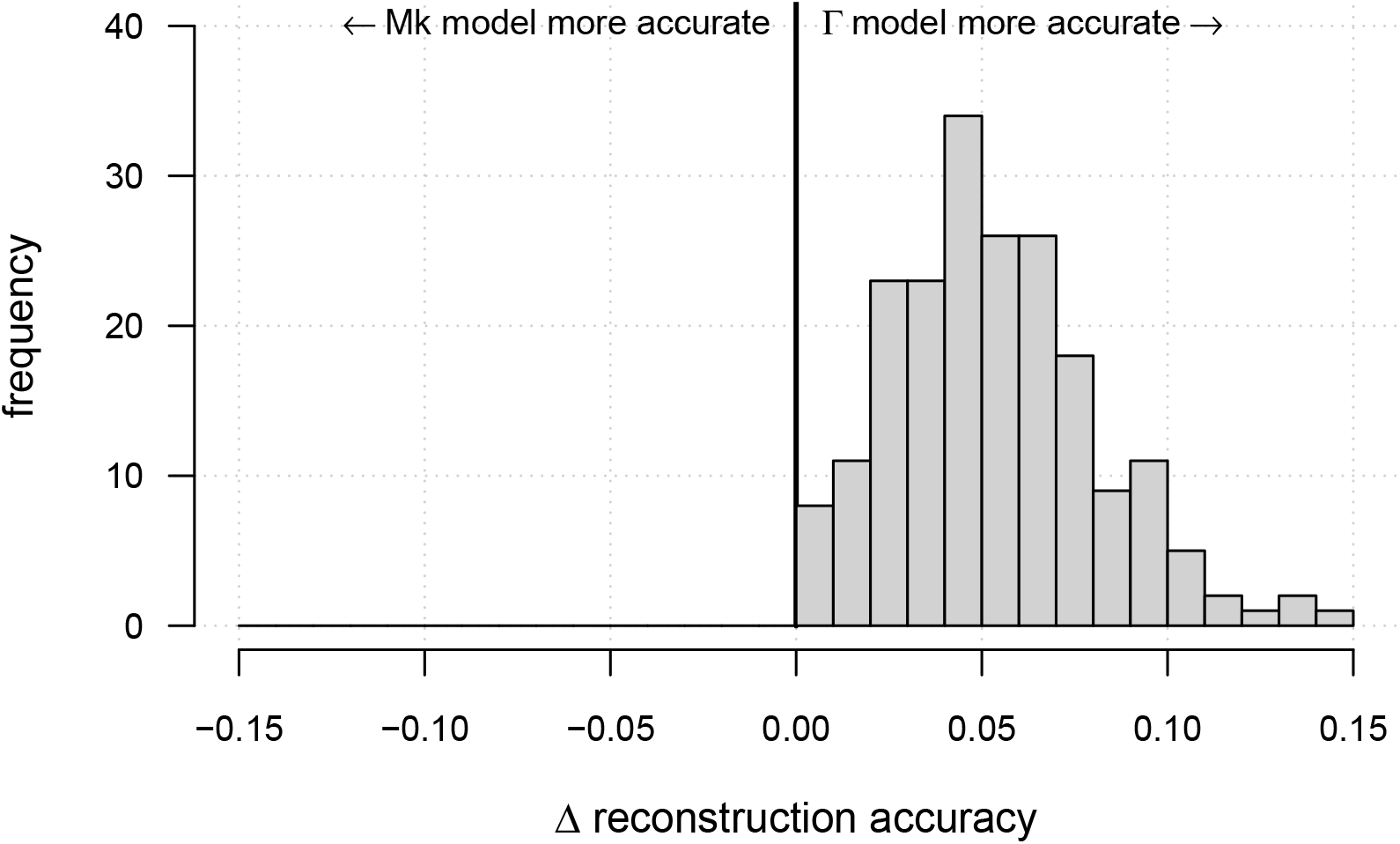
Normalized difference in accuracy of marginal ancestral state estimation under the M*k* and Γ model when the data were simulated under the Γ model. Raw accuracy was invariably higher for the generating model. See main text for more details.

### 2.5 Empirical case study

To explore the behavior of our method using empirical data we elected to analyze a dataset of forefoot and hindfoot digit number in squamate reptiles originally published by Brandley et al. (2008). These data consist of a single estimated tree and observations of 0, 1, 2, …, 5 digits in the fore and hindfeet of 257 species of squamates, plus the tuatara (*Sphenodon punctatus*). A very small number of species were reported in Brandley et al. (2008) as exhibiting polymorphism in digit number. Since the authors scored these taxa with a digit number tally equal to the *average* number of digits (and because they consist of relatively few species of the dataset), we simply rounded polymorphic taxa to the closest whole digit.

In our recently published book (Revell and Harmon 2022) we analyzed a subset of the Brandley et al. (2008) data for hindfoot digit number only and discovered that directional (sequential loss) and ordered (sequential loss and gain) M*k* models had much better support (accounting for model parameter complexity) compared to other transition models between states. As such, here we’ve elected to focus on only this category of model.

For each of the fore and hindfoot digit number traits, we fit a total of eight models: four M*k* models and four models with Γ-distributed rate heterogeneity among edges. These models were as follows (in each case, the number of parameters to be estimated is indicated in parentheses): a directional (loss-only) M*k* model with one rate of digit loss (1); a directional Γ model with one rate of digit loss (2); an ordered (loss-and-gain) M*k* model with one rate of loss and a second rate of gain (2); an ordered Γ model with one rate of loss and another rate of gain (3); a directional (loss-only) M*k* model with a different rate of loss for each character level (5); a directional Γ model with a different rate of loss for each level (6); an ordered (loss-and-gain) M*k* model with different rates of loss and gain for each character level (10); finally, an ordered Γ model with different rates of loss and gain for each level (11). As in our simulations, above, we used a FitzJohn et al. (2009) root prior, ran ten optimization iterations per model with random starting values, and set the number of rate categories of the discretized Γ distribution to *m* = 8. For the best-fitting Γ model for each trait, we also computed the marginal edge rates and scaled likelihoods (marginal ancestral states) at all nodes of the tree.

Table 1 shows a summary of the model fits, estimated value of *α* (if applicable), model parameter complexity, and model support for each of the eight trait evolution models we fit to forefoot digit number. In comparing between each *pair* of otherwise equivalent models with and without Γ rate heterogeneity among edges, information theoretic model support (measured via AIC) was consistently superior for the rate heterogeneous model (12.0 ≤ Δ*AIC* ≤ 379.9 across all model pairs; Table 1). ML estimated values of *α* across all models were invariably low (0.047 ≤ *α* ≤ 0.104), indicating relatively *high* rate heterogeneity among edges in the best-fitting Γ models. The best-supported model overall was unambiguously the ordered, multi-rate, Γ rate heterogeneous model (Table 1). Consequently, we used this model to estimate marginal edge rates of the character and scaled likelihoods (ancestral states) at all internal nodes of the phylogeny. This reconstruction is illustrated in Figure 6A.

**Table 1.** Log-likelihood, estimated value of *α* (if applicable), number of parameters, AIC, and model weights for five different discrete character evolution models of forefoot digit number in squamate reptiles, with and without Γ rate heterogeneity among edges. See main text for more details.

| | log(L) | estimated $\alpha$ | d.f. | AIC | weight |
| --- | --- | --- | --- | --- | --- |
| directional one-rate Mk | -386.5609 | – | 1 | 775.1218 | 0.00000 |
| directional one-rate $\Gamma$ | -195.6262 | 0.06039 | 2 | 395.2524 | 0.00001 |
| ordered two-rate Mk | -338.1719 | – | 2 | 680.3438 | 0.00000 |
| ordered two-rate $\Gamma$ | -192.0290 | 0.04669 | 3 | 390.0580 | 0.00013 |
| directional five-rate Mk | -197.5535 | – | 5 | 405.1069 | 0.00000 |
| directional five-rate $\Gamma$ | -190.1696 | 0.08672 | 6 | 392.3392 | 0.00004 |
| ordered ten-rate Mk | -182.0694 | – | 10 | 384.1389 | 0.00251 |
| ordered ten-rate $\Gamma$ | -175.0838 | 0.10363 | 11 | 372.1675 | 0.99731 |

**Figure 6.**
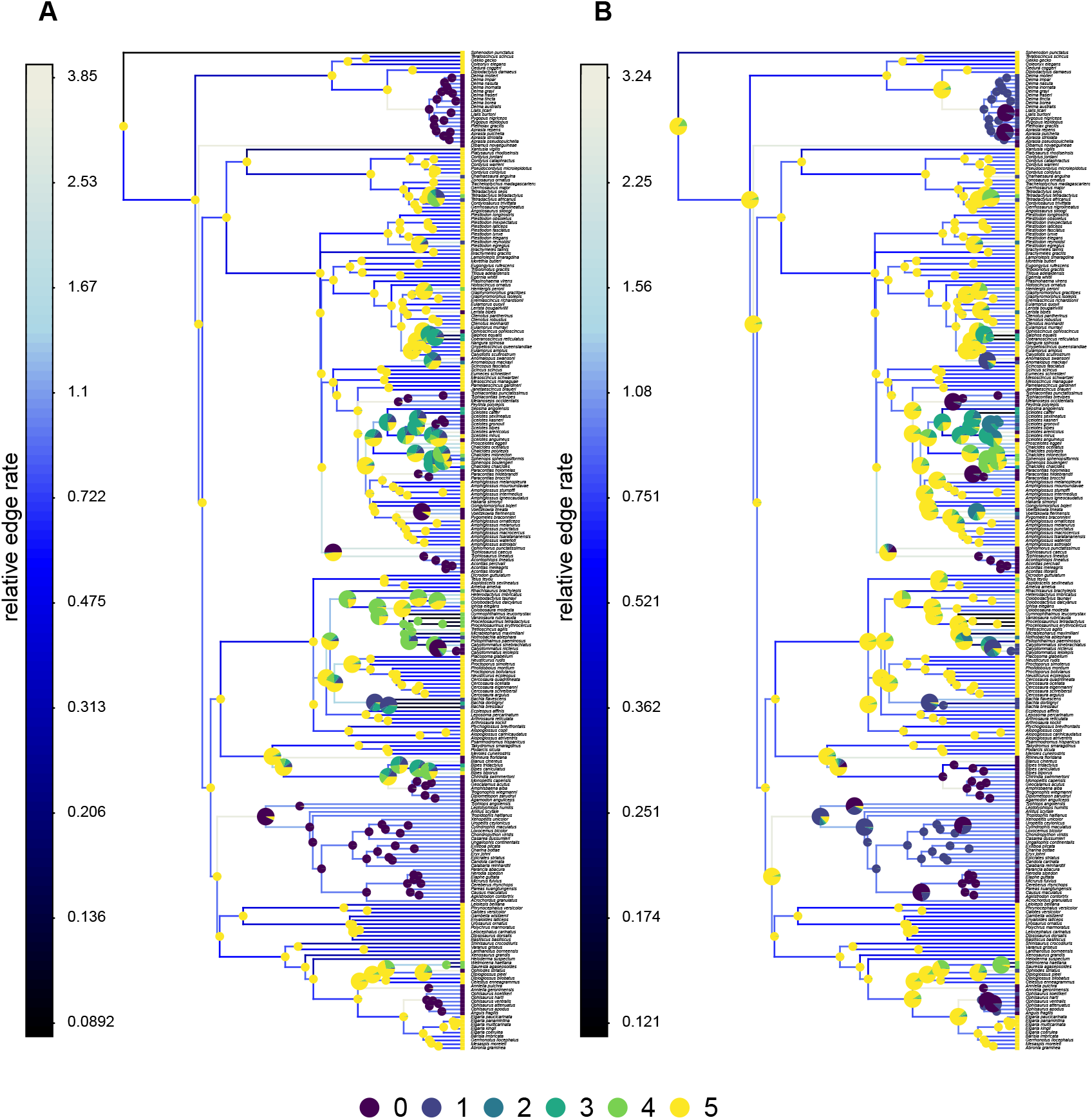
Phylogeny of squamate reptiles from Brandley et al. (2008) showing the number of digits in the forefoot (**A**) or hindfoot (**B**) of each species, estimated ancestral states under an ordered Γ model, and estimated edge rates. Larger pie charts were used for all nodes in which no state had a marginal scaled likelihood *>* 0.95. See main text for more details.

Table 2 shows a summary of the model fits, parameter complexity, and model support for each of the eight trait evolution models we fit to hindfoot digit number. Much as we found for forefoot digit number, comparing between each pair of models with and without Γ rate heterogeneity among edges, model support was consistently higher for the Γ model (14.2 ≤ Δ*AIC* ≤ 293.3 across all model pairs; Table 2). As for forefoot digit number, rate heterogeneity in the best-fitting Γ models was substantial (all 0.033 ≤ *α* ≤ 0.266). The best-supported model overall was the ordered, multi-rate, Γ rate heterogeneous model, with an Akaike weight of around 0.84. The second best-supported model was the directional, multi-rate, Γ model (Table 2). We used the best-supported ordered model to estimate marginal edge rates of the character and scaled likelihoods (ancestral states) at all internal nodes of the tree. This reconstruction is illustrated in Figure 6B.

**Table 2.** Log-likelihood, estimated value of *α* (if applicable), number of parameters, AIC, and model weights for five different discrete character evolution models of hindfoot digit number in squamate reptiles, with and without Γ rate heterogeneity among edges. See main text for more details.

| | log(L) | estimated $\alpha$ | d.f. | AIC | weight |
| --- | --- | --- | --- | --- | --- |
| directional one-rate *Mk* | -357.9915 | – | 1 | 717.9829 | 0.00000 |
| directional one-rate $\Gamma$ | -210.3328 | 0.04761 | 2 | 424.6656 | 0.00323 |
| ordered two-rate *Mk* | -324.5030 | – | 2 | 653.0060 | 0.00000 |
| ordered two-rate $\Gamma$ | -209.5280 | 0.03284 | 3 | 425.0559 | 0.00266 |
| directional five-rate *Mk* | -211.7965 | – | 5 | 433.5930 | 0.00004 |
| directional five-rate $\Gamma$ | -202.4479 | 0.26559 | 6 | 416.8958 | 0.15709 |
| ordered ten-rate *Mk* | -203.8508 | – | 10 | 427.7016 | 0.00071 |
| ordered ten-rate $\Gamma$ | -195.7757 | 0.25896 | 11 | 413.5515 | 0.83628 |

Biologically, these results tell us that there is strong evidence for rate heterogeneity in digit gain and loss across squamate reptiles. Some transitions occur more rapidly than others – which we already knew from previous analyses. This analysis gives us the additional information that particular lineages have highly elevated transition rates compared to others. There remains ambiguity as to whether digits are both gained and lost (the ordered model) or only lost (the directional model): though the former is clearly better supported for forefoot (Table 1) than in hindfoot digits (Table 2).

## 3 SOFTWARE

The model of this article is implemented in the function `fitgammaMk` of the *phytools* (Revell 2012; Revell 2024) R package. All analyses of this study were undertaken in R (R Core Team 2023) using *phytools. phytools*, in turn, depends heavily on the object classes and methods of the core R phylogenetics package, *ape* (Paradis et al. 2004; Popescu et al. 2012; Paradis and Schliep 2019). At the time of writing, *phytools* also depended on or imported from a number of other R packages including *clusterGeneration* (Qiu and Joe. 2020), *coda* (Plummer et al. 2006), *combinat* (Chasalow 2012), *doParallel* (Microsoft and Weston 2022a), *expm* (Maechler et al. 2023), *foreach* (Microsoft and Weston 2022b), *maps* (Becker et al. 2022), *MASS* (Venables and Ripley 2002), *mnormt* (Azzalini and Genz 2022), *nlme* (Pinheiro and Bates 2000; Pinheiro et al. 2022), *numDeriv* (Gilbert and Varadhan 2019), *optimParallel* (Gerber and Furrer 2019), *phangorn* (Schliep 2011), *plotrix* (Lemon 2006), and *scatterplot3d* (Ligges and Mächler 2003). The *geiger* (Harmon et al. 2008; Pennell et al. 2014) and *viridisLite* (Garnier et al. 2023) packages were also used for some analyses and visualizations of this study.

## 4 DISCUSSION

Over the past several decades, phylogenies have become pervasive in many disciplines of the biological sciences and beyond. For example, phylogenies are now employed extensively in biomedical research, in infectious disease epidemiology, in genomics, and in anthropology, among other fields (e.g., Mace and Holden 2005; Singh et al. 2009; Colijn and Gardy 2014; Brown et al. 2017; Somarelli et al. 2017). Increasingly, phylogeneticists combine the estimated tree (typically obtained from aligned nucleotide or amino acid sequences) with phenotypic trait data in an effort to learn something about how the features and attributes of living organisms have evolved over time (O’Meara 2012; Harmon 2019; Revell and Harmon 2022). For trait data that is discretely-valued, such as the presence or absence of a physical feature or a categorical behavioral trait, the typical model used to approximate character evolution on a phylogeny is one that’s been called the M*k* model (Pagel 1999; Lewis 2001; Revell and Harmon 2022; Revell 2024). The M*k* model describes a continuous-time Markov chain with *k* states, and we normally assume that this process is constant over all the branches and nodes of our reconstructed tree (but see Marazzi et al. 2012; Beaulieu et al. 2013).

In the present article we introduce a totally new model in which we assume that the edge-wise rate of trait evolution is distributed as a random variable drawn from a Γ distribution, similar to the Γ-distributed rate heterogeneity among sites model often employed in molecular phylogenetic tree inference methods (e.g., Yang 1994; Felsenstein 2004). The Γ distribution is a flexible two parameter distribution which has a general form determined by the parameter *α*. When the second parameter of the Γ distribution, *β*, is set to *β* = *α*, then the mean of the distribution is 1.0 and the variance decreases monotonically with increasing *α*. Indeed, as *α* → ∞ (but in practice for *α <<* ∞), our Γ model converges on a homogeneous rate M*k*. Conversely, as 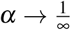 (but in practice for 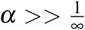) the Γ model of this paper converges on a two-rate process in which some branches evolve with a very high rate, and most not at all. There is substantial biological justification for considering models that allow rates to vary across branches in a phylogenetic tree.

When the generating process is Γ-distributed rates among edges, we found that it was possible to estimate *α* reasonably well – particularly if *α* was low and especially for larger trees (Figures 3, 4). On the other hand, we found that the variance in estimated *α* increased substantially for larger values of *α* (i.e., low rate heterogeneity) and for phylogenies with fewer tips (Figures 3, 4). We also found that the method had modest power to reject the null hypothesis of constant rates. In particular, even for relatively high rate heterogeneity (*α* = 0.5) power was only around 50% for *N* = 800. We found that taking into account rate heterogeneity when present had a highly significant impact on ancestral state estimation (Figure 5). In general, estimated ancestral states obtained for data simulated under Γ-distributed rate heterogeneity among edges were measurably more reliable when this rate heterogeneity was taken into account in the fitted model (Tables 1, 2, Figure 6).

Lastly, we analyzed a dataset of fore and rearfoot digit number in 257 species of squamate reptiles (plus the tuatara, *Sphenodon punctatus*) using a phylogeny and dataset from Brandley et al. (2008). We found significant evidence for rate heterogeneity in digit number evolution, with all Γ models fitting substantively better than their rate homogeneous counterparts.

In conclusion, we believe that the analyses we illustrate in this article can be an important addition to the growing toolbox of methodologies for studying character evolution on trees. We hope that its statistical properties, use cases, and limitations will be examined further in future research.

## 5 DATA AVAILABILITY

This article was written in Rmarkdown (Xie et al. 2018, 2020; Allaire et al. 2023), and developed with the help of both *bookdown* (Xie 2016, 2023) and the posit Rstudio IDE (RStudio Team 2020). All data and markdown code necessary to exactly rebuild the submitted version of this article (including its analyses and figures) are available at https://github.com/liamrevell/Revell-and-Harmon.fitgammaMk.

## 6 ACKNOWLEDGMENTS

This article had no direct contributors beyond the authors; however, we appreciate M. Brandley for sharing the data from his 2008 study which we have used in many workshops, as well as herein for our empirical case study.

## 7 FUNDING

This research was funded by grants from the National Science Foundation (DBI-1759940) and FONDE-CYT, Chile (1201869), to Liam J. Revell. The funders had no role in study design, data collection and analysis, decision to publish, or preparation of the manuscript.

## Notes

https://github.com/liamrevell/Revell-and-Harmon.fitgammaMk

### Competing Interest Statement

The authors have declared no competing interest.

